# Taxonomic revision of the threatened African genus *Pseudohydrosme* Engl. (Araceae), with *P. ebo*, a new, Critically Endangered species from Ebo, Cameroon

**DOI:** 10.1101/2020.10.05.326850

**Authors:** Martin Cheek, Barthelemy Tchiengue, Xander van der Burgt

**Affiliations:** Science, Royal Botanic Gardens, Kew, Richmond, Surrey, U.K.; IRAD-Herbier National Camerounais, Yaoundé, BP 1601, Cameroon

**Author notes:** Corresponding author: Martin Cheek.

**Keywords:** apomixis, Cross-Sanaga interval, disjunction, extinction, ornamental, vivipary

## Abstract

This is the first revision in nearly 130 years of the African genus *Pseudohydrosme*, formerly considered endemic to Gabon. Sister to *Anchomanes, Pseudohydrosme* is distinct from *Anchomanes* because of its 2–3-locular ovary (not unilocular), peduncle concealed by cataphylls at anthesis and far shorter than the spathe (not exposed, far exceeding the spathe), stipitate fruits and viviparous (vegetatively apomictic) roots (not sessile, roots non-viviparous). Three species, one new to science, are recognised, in two sections. Although doubt has previously been cast on the value of recognising *Pseudohydrosme buettneri*, of Gabon, it is here accepted and maintained as a distinct species in the monotypic section, *Zyganthera*. However, it is considered to be probably globally extinct. *Pseudohydrosme gabunensis*, type species of the genus, also Gabonese, is maintained in Sect. *Pseudohydrosme* together with *Pseudohydrosme ebo sp.nov*. of the Ebo Forest, Littoral, Cameroon, the first addition to the genus since the nineteenth century, and which extends the range of the genus 450 km north from Gabon, into the Cross-Sanaga biogeographic area. The discovery of *Pseudohydrosme ebo* resulted from a series of surveys for conservation management in Cameroon, and triggered this paper. All three species of *Pseudohydrosme* are morphologically characterised, their habitat and biogeography discussed, and their extinction risks are respectively assessed as Critically Endangered (Possibly Extinct), Endangered and Critically Endangered using the IUCN standard. Clearance of forest habitat for logging, followed by agriculture or urbanisation are major threats. One of the species may occur in a formally protected areas and is also cultivated widely but infrequently in Europe and the USA for its spectacular inflorescences.

## INTRODUCTION

The new species resulted in this revision was discovered as a result of a long-running survey of plants in Cameroon to support improved conservation management. The survey is led by botanists from the Royal Botanic Gardens, Kew and the National Herbarium of Cameroon-IRAD (Institute for Research in Agronomic Development), Yaoundé. This study has focussed on the Cross-Sanaga interval (*Cheek et al., 2001*) which contains the area with the highest species diversity per degree square in tropical Africa (*Barthlott et al., 1996*). The herbarium specimens collected in these surveys formed the foundations for a series of Conservation Checklists (see below). So far, over 100 new species and several new genera have been discovered and published as a result of these surveys, new protected areas have been recognised and the results of analysis are feeding into the Cameroon Important Plant Area programme (https://www.kew.org/science/our-science/projects/tropical-important-plant-areas-cameroon).

In October 2015 the last two authors found two leafless, flowering plants of a spectacular aroid in the Ebo Forest of Littoral Region, Cameroon (*van der Burgt* 1888, Fig. 1). Since these had prickles on the peduncle they were provisionally identified as *Anchomanes* Schott.

**Figure 1.**
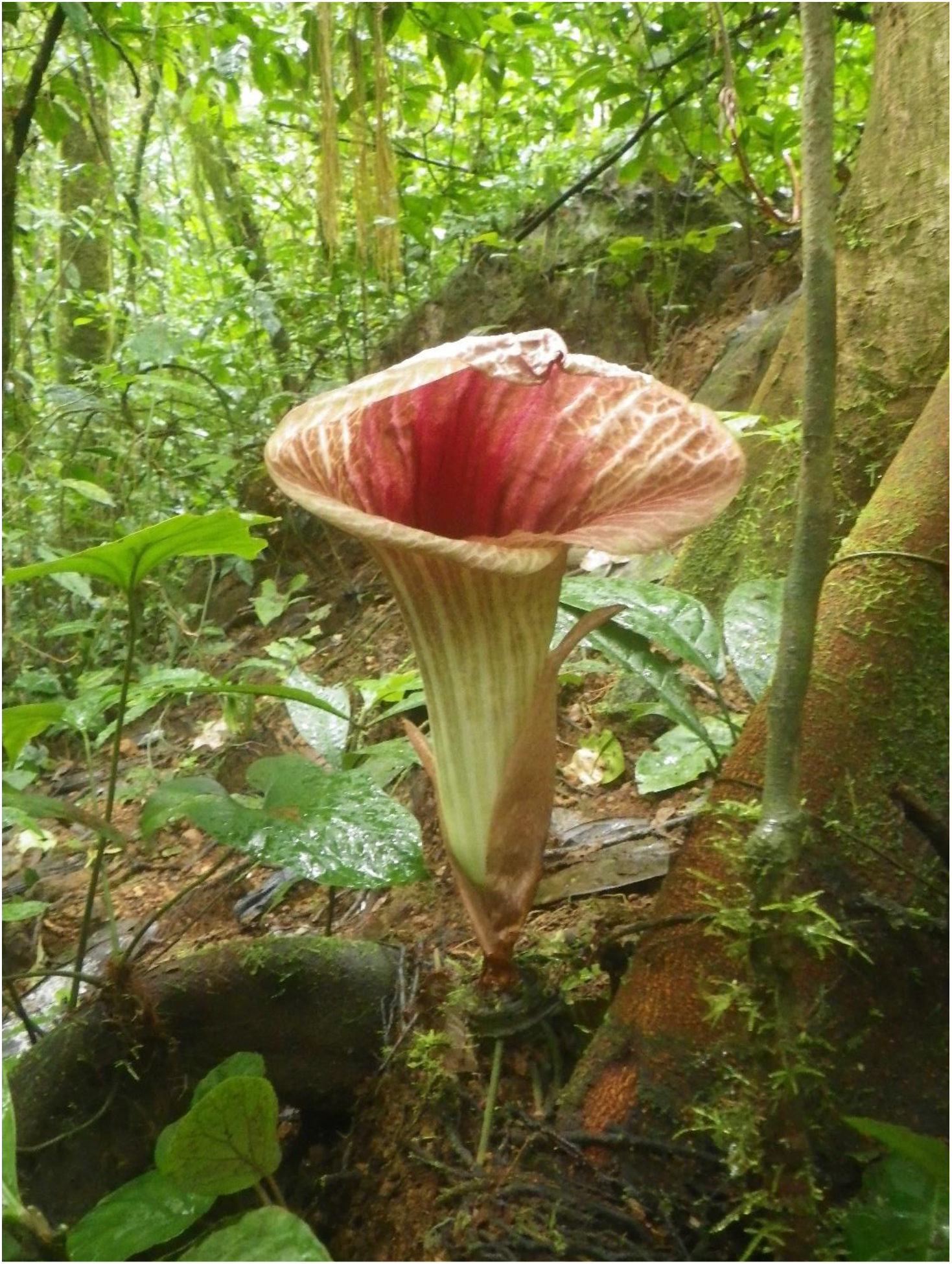
Pseudohydrosme ebo. (*van der Burgt* 1888, K, YA). Photo of flowering, leafless plant at Ebo forest, Oct. 2015. Photo by Xander van der Burgt.

In late 2018 this herbarium specimen was reidentified by the first author as *Pseudohydrosme* Engl., an erstwhile Gabonese genus previously unknown from Cameroon. *Pseudohydrosme* is separated from *Anchomanes* by a peduncle much shorter than the spathe (not far longer) and by 2–3-locular (not unilocular) ovaries (*Mayo et al. 1997*). *Van der Burgt* 1888 was suspected of representing a new species to science since it differed in several characters from the two known species of *Pseudohydrosme*. In addition, Ebo in Cameroon is separated geographically by 450 km from the range of those two previously known species in Gabon in a different biogeographic zone. In order to obtain the missing stages of fruit and leaf for *van der Burgt* 1888, it was decided to revisit the collection site at the next available opportunity. Hence, in December 2019 leaves, although not fruits, and additional field data were obtained (*van der Burgt* 2377, Fig. 2) including from a further site. Supplementary characters separating the Ebo taxon from other members of the genus were discovered in the new material. Early in 2020 the last author uncovered previously overlooked multiple additional flowering herbarium sheets, *Morgan* 25, of the same taxon at K, collected from a third site, close to the other two sites, with characters consistent with the first mentioned collection.

**Figure 2.**
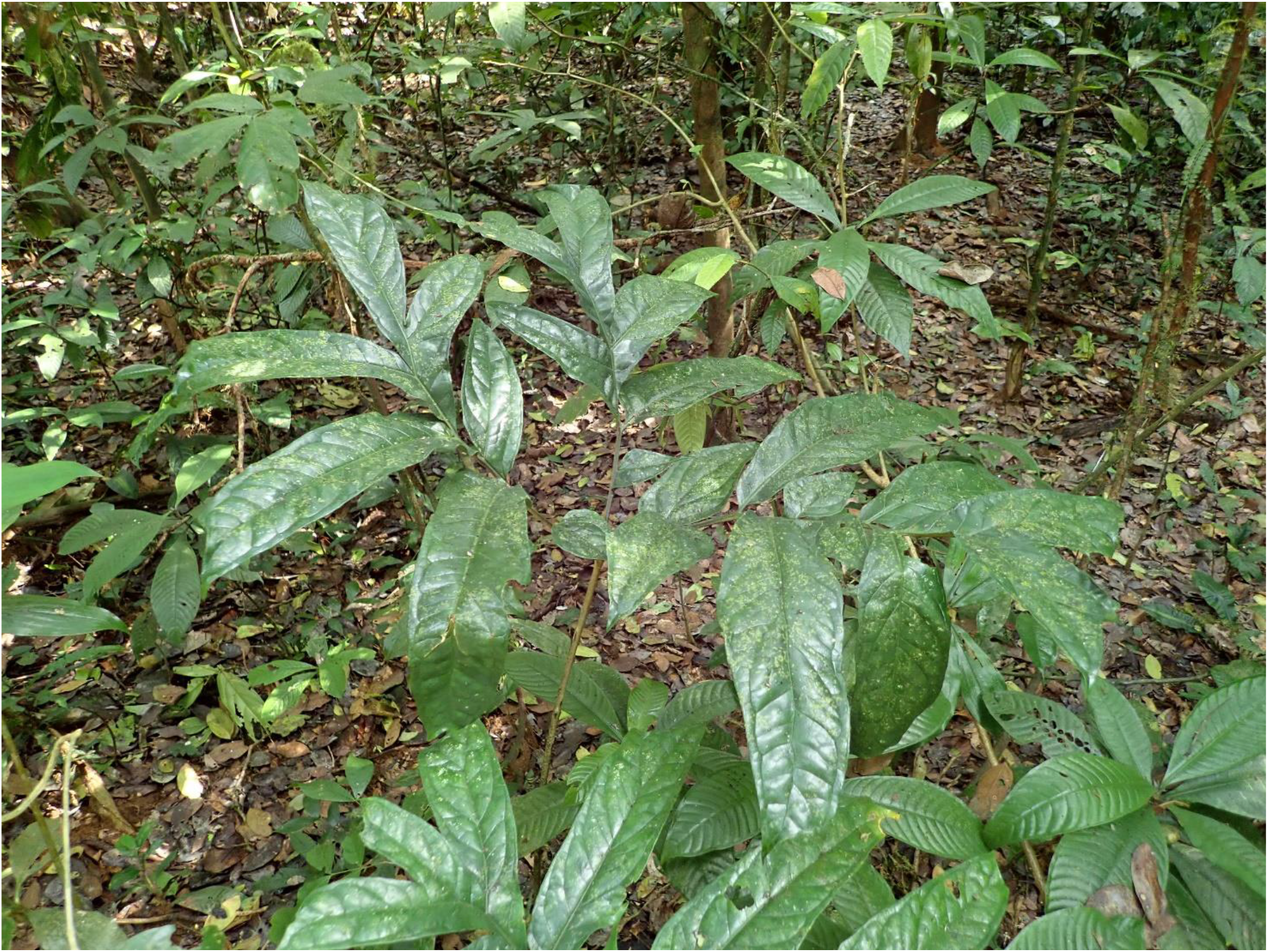
Pseudohydrosme ebo. (*van der Burgt* 2377, K, YA). Photo of plant in leaf, in habitat, Ebo forest, December, March 2019. Photo by Xander van der Burgt.

*Pseudohydrosme* was described with two species, *P. gabunensis* Engl. based on *Soyaux* 299 collected 13 Oct. 1881, and *P. buettneri* Engl. based on *Büttner* (Buettner) 519 (B) collected in Sept. 1884, both from forest at Sibang, formerly near and now largely subsumed by, Libreville, the capital and principal city of Gabon (*Engler 1892, Bogner 1981*).

In 1973 the renowned aroid specialist Josef Bogner visited Gabon and rediscovered two plants of *P. gabunensis*. Tubers were taken to Germany and cultivated allowing description of the leaves of *Pseudohydrosme* for the first time (*Bogner, 1981*).

The second species of the genus, *P. buettneri* has never been refound. It differs so greatly from the first in the structure of its inflorescence (see key and species account below) that the noted aroid specialist N.E. Brown erected a separate genus, *Zyganthera* N.E. Brown for it. *Engler* (*1911*) reduced *Zyganthera* to sectional rank within *Pseudohydrosme*.

In their monumental account ‘The Genera of Araceae’, *Mayo et al*. (1997) placed *Pseudohydrosme* next to *Anchomanes*, together with *Nephthytis* Schott in the tribe Nephthytideae Engl. Molecular phylogenetic analysis has subsequently supported the sister relationship of *Pseudohydrosme* and *Anchomanes*, but since each were represented by only a single taxon, it was not possible to test their monophyly (*Cabrera et al., 2008*; *Nauheimer et al.2010, Cusimano et al. 2011*). In presenting new data on *Pseudohydrosme gabunensis* based on successful pollination of flowers and fruit production on plants in cultivation in the Netherlands, *Hetterscheid & Bogner* (2014) questioned the distinction of *Pseudohydrosme* and *Anchomanes*. They considered the only difference to be the locularity of the ovaries (2–3 versus 1) and set aside the difference in peduncle: spathe proportions maintained in *Mayo et al*. (*1997*). However, later in their paper *Hetterscheid & Bogner* (*2014*) then brought to light two new synapomorphies that further support the distinction of *Pseudohydrosme* from both *Anchomanes* and from all other aroids (see discussion below). *Hetterscheid & Bogner* (*2014*) made the case to extend the range of *Pseudohydrosme* from Gabon southwards into the Republic of Congo (Congo-Brazzaville) citing photos and a specimen deposited at WAG by Ralf Becker. However, the species concerned is not identified and we have not been able to access the material in order to verify this statement (see methods).

In this paper we describe the Cameroon material from Ebo Forest as a new species to science, *Pseudohydrosme ebo* Cheek, in the context of a revision of the genus, last revised nearly 130 years ago (*Engler 1892*).

## MATERIALS & METHODS

Herbarium citations follow Index Herbariorum (*Thiers et al., 2020*). Specimens were viewed at B, BR, K, P, WAG, and YA. *Pseudohydrosme* is centred in Gabon. The national herbarium of Gabon is LBV, but the most comprehensive herbaria for herbarium specimens of that country are P and WAG. The National Herbarium of Cameroon, YA, was also searched for additional material but without success. During the time that this paper was researched, it was not possible to obtain physical access to material at WAG (due to the transfer of WAG to Naturalis, Leiden, subsequent construction work, and covid-19 travel and access restrictions). However images for WAG specimens were studied at https://bioportal.naturalis.nl/?language=en and those from P at https://science.mnhn.fr/institution/mnhn/collection/p/item/search/form?lang=en_US. We also searched *JStor Global Plants* (*2020*) for additional type material of the genus, and finally the Global Biodiversity Facility (GBIF, www.gbif.org accessed 23 Aug 2020) which lists 28 occurrences and 21 images, mainly relating to the holdings (including duplicate herbarium sheets) of WAG, followed by P.

Binomial authorities follow the International Plant Names Index (*IPNI, 2020*). The conservation assessment was made using the categories and criteria of *IUCN* (*2012*). GeoCAT was used to calculate red list metrics (*Bachman et al., 2011*). Spirit preserved material was not available. Herbarium material was examined with a Leica Wild M8 dissecting binocular microscope fitted with an eyepiece graticule measuring in units of 0.025 mm at maximum magnification. The drawing was made with the same equipment using Leica 308700 camera lucida attachment. The herbarium specimens of the new species described below as *Pseudohydrosme ebo* were soaked in warm water to enable the spathe to be folded back, exposing the spadix and hydrated flowers. The terms and format of the description follow the conventions of *Mayo et al*. (*1997*). Georeferences for specimens lacking latitude and longitude were obtained using Google Earth (https://www.google.com/intl/en_uk/earth/versions/). The map was made using SimpleMappr (https://www.simplemappr.net).

## RESULTS

### TAXONOMIC TREATMENT

*Pseudohydrosme* is sister to *Anchomanes* Schott (7–8 species) and the pair are in a sister relationship with *Nephthytis* Schott (6 species). Together these three tropical African (all are absent from Madagascar, but anomalous *Nephthytis bintuluensis* A.Hay, Bogner & P.C. Boyce occurs in Borneo (*Hay et al,. 1994; Nauheimer et al., 2012*)) genera comprise the Nephthytideae which is sister to the bigeneric SE Asian group Aglaonemateae Engl. (*Cabrera et al., 2008*). Both groups share adjacent male and female flower zones, free stamens, and collenchyma arranged in threads peripheral to the vascular strands of leaf blades and petioles (with the exception of *Nephthytis*, in which collenchyma can form interrupted bands (*Keating, 2002*; *Cabrera et al., 2008*). Morphological support for Nephthytideae is the primary leaf venation with basal ribs of the primary veins very well developed, i.e., ± tripartite primary development (*Cabrera et al. 2008*) Dracontioid leaf divisions characterise *Pseudohydrosme* and *Anchomanes* but are not present in *Nephthytis*. Differences between *Pseudohydrosme* and *Anchomanes* are re-assessed in the discussion below.

***Pseudohydrosme*** Engl. (*Engler 1892*: 455; Brown in *Thistleton-Dyer 1901*: 160; *Engler 1911*: 47; *Mayo et al., 1997*: 221–222)

Type species: *Pseudohydrosme gabunensis* Engl. (Lectotypified by N.E.Brown in *Thistleton-Dyer 1901*:160).

*Zyganthera* N.E. Br. (Brown in *Thistleton-Dyer, 1901*:160). Heterotypic synonym.

Large, seasonally dormant, monoecious herbs. Rhizome shallowly subterranean, the growing point at ground level, subglobose or cylindrical with annular leaf scars, and erect to horizontal or obliquely inclined, growing continuously and not renewed with each growing period. Roots fleshy, produced along length of rhizome, sometimes reproductive (the distal ends rising to the surface and producing new plants). Leaf solitary, large; petiole cylindrical, erect, long, with minute and sparse prickles, sheath very short and inconspicuous. Blade transitioning from simple, sagittate and entire in seedlings, older plants developing slits and divisions, in mature plants leaves dracontoid : trisect, primary divisions pinnatisect, distal lobes mostly truncate and bifid, sessile and decurrent, proximal lobes acuminate; primary lateral veins of ultimate lobes pinnate, forming a regular submarginal collective vein (*P. ebo*) or an irregular collective vein, or veins running into margin (*P. gabunensis*), higher order venation reticulate.

Inflorescence solitary, appearing separately from the leaf. Cataphylls papery-membranous, (3–)4–6, proximal ± triangular, small, distal oblong-elliptic, concealing spathe tube. Peduncle concealed by cataphylls at anthesis, terete, very short <1/10th the length of the spathe, with minute, sparse, prickles. Spathe large, fornicate, resembling the horn of a euphonium, unconstricted, blade very broad, with flaring auriculate margins; tube convolute, fleshy, obconic, with a few sparse prickles on the outer surface proximally. Spadix short, about 1/10–1/4 length of spathe, sessile, female zone subcylindric, male zone cylindric, obtuse or rounded, subequal to ± twice (±four times in *P. buettneri*) as long as female, completely covered in flowers and fertile to apex (*P. gabunensis*) or with a distal appendix twice as long as the fertile portion and covered in sterile male flowers (*P. buettneri*) or flowers covering only about 50% of axis in the female zone (*P. ebo*).

Flowers unisexual, perigone absent. Male flower 2–5-androus, stamens free, subprismatic, compressed, anthers sessile, connective thick, broad, overtopping thecae, thecae oblong, long, lateral, dehiscing by apical pore. Pollen extruded in strands, inaperturate, ellipsoid-oblong, very large (mean 106 micrometres diam.) exine psilate to slightly scabrous. Sterile male flowers (*P. buettneri*) composed of subprismatic, free staminodes. Female flower ovary globose to broadly ellipsoid, usually prismatic, 2–3-locular, ovules 1 per locule, anatropous, funicle short, placentation axile, at base of septum, stylar region attenuate to cylindric, narrower than ovary, stigma thick, shallowly 2–3-lobed or subdiscoid, concave centrally, wet when receptive.

Berry white, ripening dark purple, fleshy, wrinkled when mature, oblong-ellipsoid, laterally compressed to slightly bilobed, stipitate, large, borne on a slightly accrescent peduncle, stigma and style persistent (known only in *P. gabunensis*). Seeds subglobose to broadly ovoid, one side convex, the other slightly flattened, testa thin, whitish, smooth, papery, transparent; embryo large, outer surface green, inner white, raphe distinct, hilum and micropyle purple, plumule with leaf primordia. Three species.

**Phenology:** flowering Sept. and Oct. (or March in cultivation in Europe); in leaf Dec.-April. **Distribution & habitat:** Cameroon and Gabon, lowland evergreen forest on coastal sediments (Gabon) or inland foothills on basement complex rocks (Cameroon) (Fig. 3).

**Figure 3.**
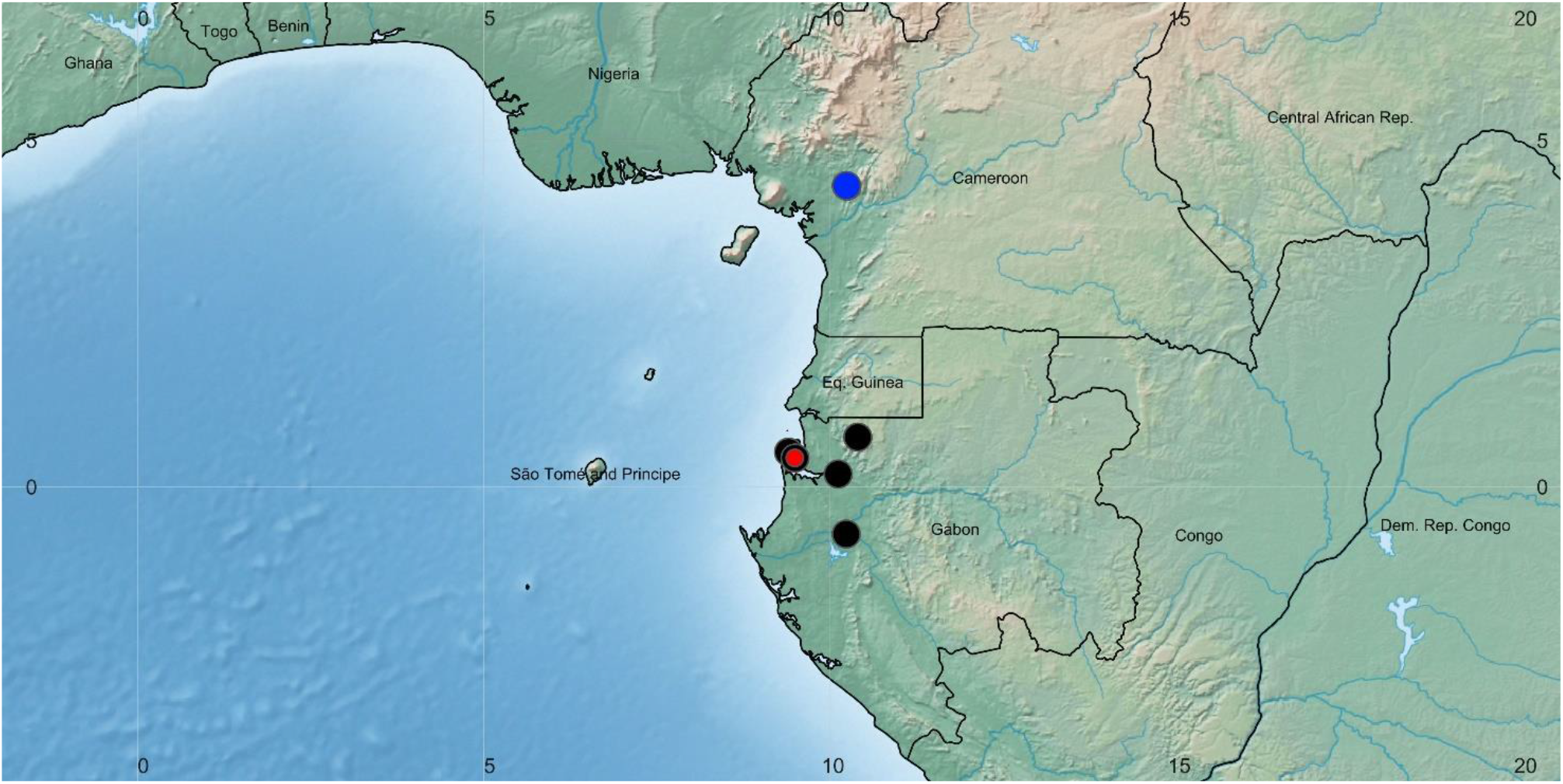
Global distribution of the species of the African genus *Pseudohydrosme*. Red dot = *P. buettneri*; black dots= *P. gabunensis;* blue dot = *P. ebo*.

**Etymology:** meaning “false *Hydrosme”, Hydrosme* Schott now being a synonym of *Amorphophallus*.

**Local name and uses:** none are documented.

**Conservation:** all species are highly infrequent and globally threatened according to IUCN (2012) criteria (see species accounts below), and *P. buettneri* is possibly extinct (not seen for over 100 years, the majority of its former habitat destroyed).

Pollination has not been investigated in detail in *Pseudohydrosme*,but is almost certainly by insects as is usual in Araceae. Two different species of flies, and two of beetles were reported to visit *P. gabunensis* (see below). In cultivation the stigmas are reported to be wet and receptive for only two days, and the scent reported to be faint, of lettuce (*Lactuca*) in the same species (see below also).

Following successful fertilisation, seed development is reported to take up to 10 months in *P. gabunensis* (see below). Seed dispersal is probably by either ground-dwelling mammals or birds consuming the thinly fleshy purple berries.

DNA analysis was performed by *Cabrera et al*. (*2008*) for *Pseudohydrosme gabunensis*, using five regions of coding (*rbcL, matK*) and noncoding plastid DNA (partial *trnK* intron, *trnL* intron, *trnL* – *trnF* spacer). The voucher was *Wieringa 3308* (WAG), GenBank codes are AM905760, AM920582, AM932319 + AM933315.

Cultivation of one species, *Pseudohydrosme gabunensis* is widespread but infrequent outside of Africa in the tropical glass-house collections of several large botanical gardens, mainly in Europe and N. America (see under that species).

Chromosome numbers are reported of one species, *Pseudohydrosme gabunensis, as* 2n = ca. 40 (*Mayo et al., 1997*); x = 13 ((*Bogner & Petersen, 2007*; *Cusimano et al., 2011*).

Germination in *Pseudohydrosme gabunensis* is cryptocotylar and takes 3 weeks to 10 months. The large seed embryo remains buried, producing a single hastate seedling leaf (*Hetterscheid & Bogner, 2014*).

Medicinal uses, and chemistry is unreported in *Pseudohydrosme*. However, the much more frequent sister genus *Anchomanes*, is harvested as a traditional medicine e.g. in Cameroon, and contains bioactive compounds (*Cheek, 1992*)

### Identification key to the sections and species of *Pseudohydrosme*

1. Spadix with distal half covered in sterile flowers. **Sect. *Zyganthera ………… P. buettneri***
1. Spadix lacking sterile flowers, distal part with male flowers only. **Sect. *Pseudohydrosm*e *…………*** 2
2. Leaf veins running to margin, not forming a regular submarginal collective vein; male and female areas of spadix contiguous, entire spadix covered in flowers; spathe blade inner surface yellow, greenish yellow or white with abrupt transition to a central dark red area; stigmas 2(–3)-lobed. Gabon (possibly Congo) ***…………* 2. *P. gabunensis***
2. Leaf veins forming a conspicuous, regular submarginal collective vein: male and female areas of spadix incompletely contiguous, female zone with axis partly naked, spathe blade inner surface light reddish brown or pink, with wide green veins, very gradually becoming darker towards the centre; stigmas 3(–2)-lobed. Cameroon ***…………* 3. *P. ebo***

Sect. ***Zyganthera*** (N.E.Br.) Engl. (*Engler 1911:* 49)

Type (and only) species: *Pseudohydrosme buettneri* Engl.

*Zyganthera* N.E. Br. (Brown in *Thistleton-Dyer, 1901*:160). Homotypic synonym

Male flowers connate in pairs; distal half of spadix covered in sterile male flowers; ratio of female:male (including sterile male) spadix portions c.1:4

**1. *Pseudohydrosme buettneri*** Engl. (*Engler 1892*:456; *Engler 1897*: 59; Brown in *Thistleton-Dyer 1901*: 160; *Engler 1911*: 49). – Fig. 3, 4.

**Figure 4.**
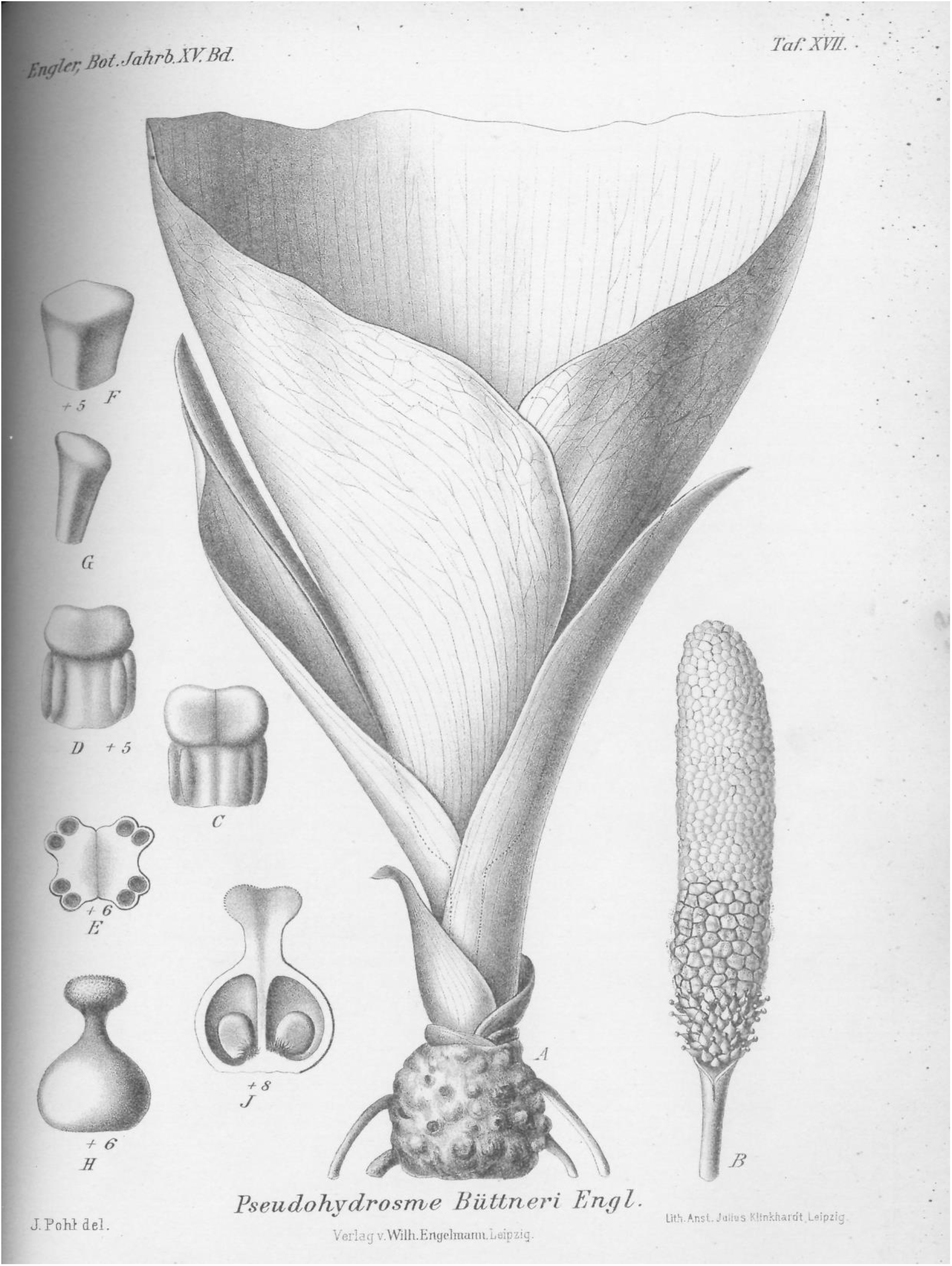
Pseudohydrosme buettneri. (*Buettner* 519, B). Drawing of flowering plant. (A) habit, rhizome and inflorescences; (B) spadix after removal of spathe showing from base to apex, female, male and staminodal flowers; (C & D) paired male flowers, side view; (E) paired male flowers, transverse section; (F & G) staminodes, sterile male flower; (H) female flower, entire, side view; (J) female flower, longitudinal section. Reproduced from protologue, (*Engler, 1892*: taf. XVII). Drawn by Josef Pohl.

Holotype: Gabon, Estuaire Province, Libreville “Gabun, Mundagebiet; Sibange-Farm” fl. Sept. 1884, *Buettner* 519 (Holotype B destroyed or mislaid).

Terrestrial herb, rhizome vertical, subglobose, 2.5 cm long, 2.5 cm wide, surface tuberculate, roots fleshy, from along the length of the rhizome. Leaf unknown.

Inflorescence: Cataphylls three or more, 1.75–6 × 0.9–1 cm; peduncle 3 cm long, colour and indumentum unknown. Spathe 80 cm long. Spadix subcylindrical, 7–8 cm long, c. 1.3 cm diam. Female zone 1.3 cm long. Fertile male zone 2 cm long. Appendix (sterile male flowers) 5 cm long, c. 1.5 cm diam.

Female flowers 4 mm long, ovary globose, 3 mm diam., style 1 mm long, slender; stigma 1 mm diam., thick. Male flowers. Stamens 2 mm long, 2 mm wide, usually the two stamens of a flower close together. Staminodes prismatic, 4–6 sided, lacking anther thecae and much smaller in diameter than the stamens (from *Engler* (*1892*:456 and fig. XV).

**Phenology:** flowering in September.

**Local name and uses**: none are known.

**Etymology:** named for the collector of the only known, and type specimen, Oscar Alexander Richard Buettner (Büttner) – (1858–1927), traveller and collector.

**Distribution & ecology:** known only from Sibang in Libreville, coastal lowland evergreen forest dominated by *Aucoumea klaineana* Pierre.

**Additional specimens:** none are known.

**Notes:** *Pseudohydrosme buettneri* has the largest inflorescence by far of all known species of the genus, with an 80 cm long spathe. The type specimen had lost the top part of the spathe, but dimensions were given by the collector (*Engler, 1892*).

The type, and only known specimen was at B, but is reported to be no longer there (Bogner 1981). One can deduce that it was destroyed in the allied bombing of Berlin in March 1943, when most of the specimens at B were also destroyed, however the type specimen of *P. gabunensis* (see below) dating from about the same time, and also housed at B, has survived.

No additional specimens of this species have been found in the 136 years ensuing from collection of the type specimen. *Hetterscheid & Bogner* (*2014*) have questioned whether this species is not just a variant of *P. gabunensis*. However, this seems highly unlikely, because the specimen differs in three independent characters from *P. gabunensis* (and *P. ebo*):

1. the ratio between the female zone and the male zone (of fertile and sterile flowers) differs greatly between the two. In *P. buettneri* the ratio is 1:4+, while in the other two species it is less than 1:2.
2. in *P. buettneri* most of the spadix consists of an appendix of sterile male flowers. No such sterile appendix occurs in the other two species.
3. in *P. buettneri* the stamens are paired (*Engler 1892*), while in the other two species the stamens are not paired, but present in an indistinct ring of five.

Additional differences between the species seen in the pistils and spadices. The styles in *P. buettneri* are <1/4 the width of the ovary. In *P. gabunensis* it is ½. The spadix of *P. buettneri* is cylindrical, and even in width along its length, while that of *P. gabunensis* shows a pronounced constriction at the junction of female and male zones, and the male zone reaches a greater width than the female zone.

**Conservation**. *Pseudohydrosme buettneri* is here assessed as Critically Endangered (Possibly Extinct). This is because it has only been found once, at a single site, in the “Munda region” at Sibang Farm or Plantation, in 1884. At that time Sibang was far outside Libreville consisting largely of forest, some of which was exploited to produce forest products such as timber and rubber, and cleared to produce agricultural products by Europeans for international commerce e.g. by the Woermann company (*Cheek et al. 2011*: 45). The Munda is the estuary that forms the eastern edge of the peninsula on which Libreville sits. Tributaries of the Munda drain the Sibang area. Beginning in 1960, the population of Libreville expanded 200-fold, and its footprint expanded. Only a small part of the original forest formerly known as Sibang survived. This part measures about 400 m x 400 m as measured on Google Earth (see further details under *P. gabunensis*, below) and is now entitled the ‘Sibang Arboretum’. This minute remnant of forest is probably the most visited by botanists in the whole of Gabon because it is immediately adjacent to the site of the current National Herbarium, LBV (Cheek, pers. obs.). In the unlikely although hoped-for rediscovery of *Pseudohydrosme buettneri*, the area of occupancy would be expected to be calculated as 4 km^2^ using the IUCN preferred gridcells of this size, and the extent of occurrence of the same size. If it should be found anywhere in the vicinity of Libreville it is likely to be threatened by human pressures since most of the population of Gabon is concentrated here.

The Libreville region has the highest botanical specimen collection density in Gabon, with 5359 specimens recorded in digital format. It also the highest level of diversity of both plant species overall and of endemics (Sosef 2005). The coastal forests of the Libreville area are known to be especially rich in globally restricted species (*Lachenaud et al. 2013)*. These authors detail 19 species globally restricted to the Libreville area, of which eight have not been seen recently and which are possibly extinct. Among these is *Octoknema klaineana* Pierre, a rainforest tree “only collected in the immediate area of Libreville at the beginning of the 20^th^ century, and only once since.” (*Gosline & Malecot, 2011)*. Most of the collections of this possibly extinct species of *Octoknema* were also, as with *Pseudohydrosme buettneri*, from Libreville-Sibang, and were mainly made in the period 1896–1912, during the colonial period, before the city expanded to its current extent. The other seven species recorded as globally restricted to the Libreville area and as possibly extinct by *Lachenaud et al*. (*2013*): are *Ardisia pierreana* Taton (*Taton,1979*), *Dinklageella villiersii* Szlach. & Olszewski (*Szlachetko & Olszewski, 2001*), *Eugenia librevillensis* Amshoff (*Amshoff, 1958*), *Hunteria hexaloba* (Pichon) Omino (*Omino,1996*), *Pandanus parvicentralis* Huynh (*Huynh, 1986*), *Psychotria gaboonensis* Ruhsam (*Ruhsam et al. 2008*) and *Tristemma vestitum* Jacq.-Fél. (*Jacques-Félix, 1986*). These species have also not been seen in several decades, or more, in the case of the penultimate species, since 1861. The explanation for this hotspot of unique species, fast disappearing if not already extinct, at Libreville may be that it has the highest rainfall in Gabon (*Gosline & Malecot, 2011*), with c.2.9 m p.a.

It seems likely that *Pseudohydrosme buettneri* is an additional lost endemic species to the Libreville area, likely rendered extinct by the expansion of the city. Let us hope it is rediscovered in a fragment of forest in the greater Libreville area, although this seems extremely unlikely given that it was the most spectacular species of the genus with by the largest spathe (80 cm long) known in the genus, and that as stated above, the Libreville area is the most intensively botanically surveyed part of Gabon (*Sosef et al., 2005)*.

Sect. ***Pseudohydrosme***

Type species: *Pseudohydrosme gabunensis* Engl.

*Chorianthera* Engl. (*Engler, 1911*: 48). Homotypic synonym

Male flowers free, in clusters of c. 5; distal half of spadix lacking sterile male flowers; ratio of female:male spadix portions 1:2

***2. Pseudohydrosme gabunensis*** Engl. (*Engler, 1892*:455; *Engler 1897*:59; *Brown, 1907*:161; *Engler, 1911*:48; *Bogner, 1981*:33; *Hetterscheid & Bogner, 2014*:104–113). – Fig. 3, 5.

**Figure 5.**
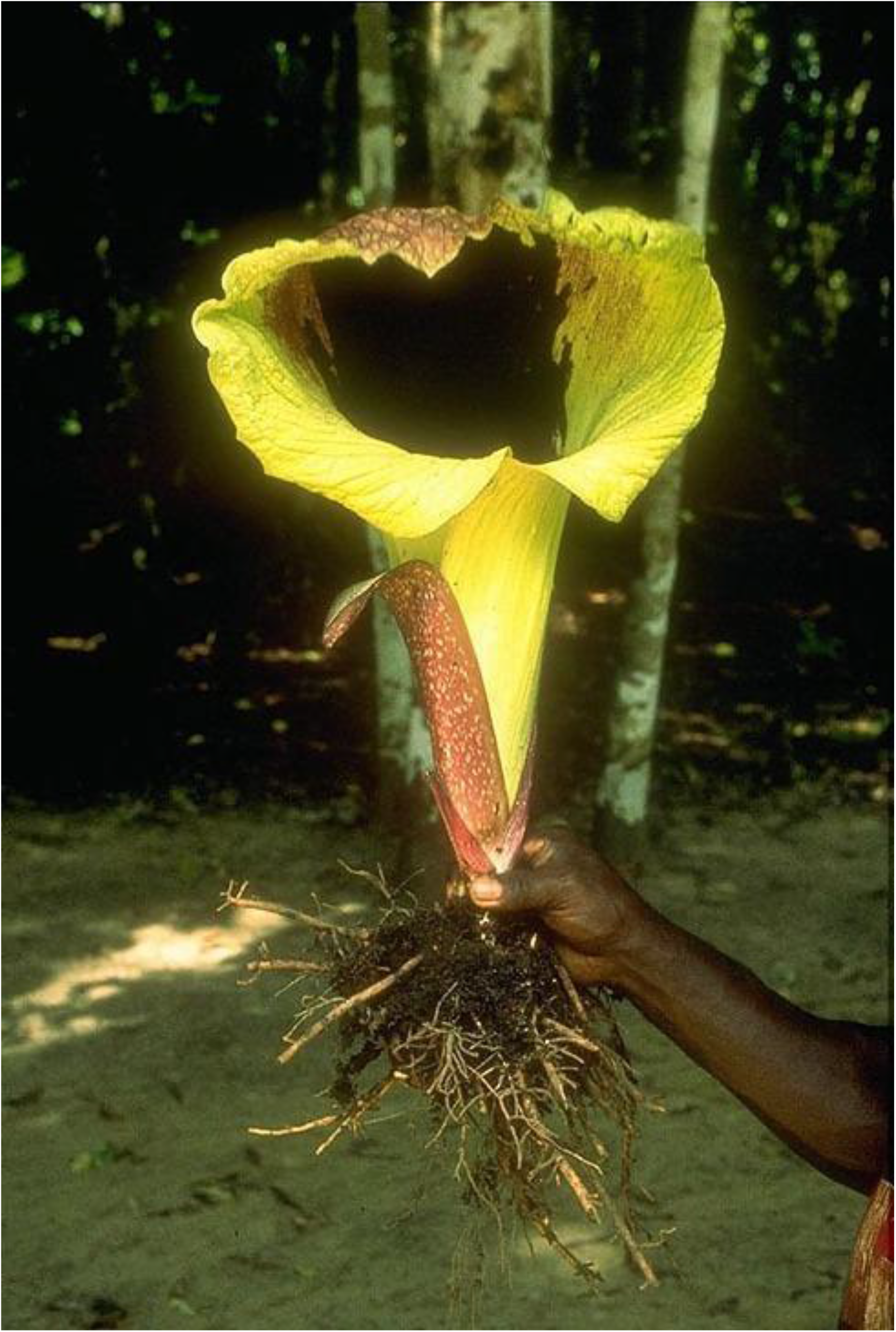
Pseudohydrosme gabunensis. *(Bogner* 664, K, M, US). Photo of flowering, leafless plant at Sibang, Libreville, Gabon, 29 Oct. 1973. Photo by Josef Bogner/License: Creative Commons (cc-by-sa-nc-3.0)

**Holotype:** Gabon, Estuaire Province, Libreville, Sibang, “Gabun, Mundagebiet; Sibang-Farm am Ufer des Maveli” fl. 13 October 1881, *Soyaux* 299 (Holotype: B100165306, Herbarium specimen, image!)

Terrestrial herb, rhizome light brown, ellipsoid or subcylindric, erect or oblique, 9–12 cm diam. to 15 cm long, surface with transverse ridges. Roots fleshy 5–8 mm thick, brownish yellow, sometimes developing new plants at their tips.

Leaf 1–1.3(–2.2) m tall, petiole terete, 1–1.4 cm diam. at base, dark green olive and spotted with small yellowish white points; spines 1–2 mm long. Blade of youngest seedlings sagittate-elliptic c. 5 cm long, 3–4 cm wide, basal sinus c. 2 cm long, breadth variable (see *Hetterscheid & Bogner, 2014*). Successively formed blades developing slits and divisions. Blade of mature leaves dracontoid, primary divisions 30–35 cm long, pinnatisect, lobes 5–8, dimorphic, larger, distal lobes elliptic (4–)8–23 cm long, (2–)3–7(–11) cm, apex truncate, bifid, (0.5–)1–3 cm long; smaller, proximal leaflets ovate, 4.5–8 cm long, 2.5–5 cm wide, apex cuspidate; lateral nerves 4–8 on each side of the midrib, conspicuous on abaxial surface, running to the margin or forming an incomplete submarginal nerve, higher order nerves reticulate.

Inflorescence: Cataphylls 4–6 membranous, reddish white or brown-purple, slightly spotted distichous, proximal subtriangular shorter, distal becoming longer and oblong elliptic to the spathe 1.5–29 cm long, (1–)2–2.5 cm wide; peduncle (3–)5–9 cm long, 1–1.5 cm diam., with minute sparse spines 1–2 mm long and greenish white, colour as petiole. Spathe (30–)40–55 cm long, basal half (20–25 cm long) funnel-shaped to tubular, fleshy and to 5 mm thick, blade comprising the distal half of the spathe, flaring widely and curving forward, the apex obtuse, margin undulate.

Outer surface uniform bright pale yellow, greenish yellow or yellow white. Inner surface of blade pale yellow or yellowish white, at marginal separated by a solid irregular line from the dark purple central area which continues to the base of the tube. Mouth facing horizontally, usually orbicular or elliptic. Spadix with “unpleasant smell, but not so strong as some Araceae” (van der Laan 7641, WAG), sessile, cylindrical, (6–)9–12.5 cm long, (1.5–)2–2.5 cm diam. Female zone (2–)3.5(–4) cm long, female flowers completely covering the surface of the axis, usually contiguous with the male zone. Male zone (3.5–)6–8.5 cm long, apex rounded, completely covered in fertile male flowers. Sterile appendix absent.

Flowers lacking perigone. Male flowers with c. 5 stamens, stamens densely packed, free, sessile, 4 mm long, in plan view isodiametric, subprismatic, 5–6-faceted, in cross section c. 1.8(–2) mm x 1.2 mm wide, apex convex purple, sides white, anther thecae c. 3mm long, opening by an apical pore, pollen orange or yellow, in strings. Female flowers white with ovary yellowish-white globose or ellipsoid, 2–3 mm diam., 2(–3)-celled; style 1–1.5 mm long, 1.5 mm diam., stigma black to reddish brown, surface papillose, 2 mm wide, bilobed, lobes with a broad concave area, apex rounded.

Berry white, ripening purple-black, surface wrinkled when ripe, thinly fleshy, transversely ellipsoid, laterally compressed, rarely globose, 0.8–1.2 cm long, 1.5–1.6 cm wide, style and stigma persistent, (1–)2-seeded, apex rounded-truncate, base stipitate stipes (2–)3–4 mm long, c. 2 mm diam. Seeds subglobose to broadly ellipsoid, one side flattened, the other convex, 9 mm long, 7 mm wide.

**Phenology:** flowering mid-Sept.-late Oct.

**Distribution & ecology:** Gabon, Estuaire, Moyen-Ogooué (probably) and Woleu-Ntem Provinces, known from five sites in lowland rainforest sometimes with *Aucoumea gabonensis* Pierre (Burseraceae); 0–100 m alt.

**Etymology:** meaning “coming from Gabon” (formerly, in German “Gabun”).

**Local names & uses:** none known.

**Additional specimens: Gabon, Woleu-Ntem Province**, c. 15 km NE Asok, 600–700 m alt., (leg. *Breteler & de Wilde* s.n. 21 Aug. 1978) cult. Wageningen, fl. 13 March 1984, *van der Laan* 764 (Bot. Gard. No. 978PTGA550), WAG0351246, WAG0351247 images!); **Estuaire Province, Libreville, Sibang:** “Sibang, hinter der Station forêstier; wächst im sandigen Lehmboden im Regenwald, sehr schattig, c. 20 m, fl. 29 Ocktober 1973, *Bogner* 664 (K!, M n.v. US n.v.); Sibang, st. 10 April 1994, *Wieringa & Haegens* 2710 (WAG0181636, WAG0181637 images!); Sibang Forest, st. 1 Dec. 1994, *Wieringa* 3308 (WAG0181631, WAG0181632, WAG 0181633, WAG0181634, WAG0181635 images!); Sibang Arboretum fl. 25 Oct. 2005, *Sosef et al*. 2029 (WAG 0223594, WAG0223595 images!, WAG8004057, WAG0108030, WAG.1665445); **Kango**, plantations de Assouko, près de poste de Kango, le Komo (estimated as 0° 10’ 41.8” N, 10° 06’ 45.54”E), fl. 2 Oct. 1912, *Chevalier* 26828 (P02093245 image!); **Forêt de la Mondah**, road from Libreville to Santa Clara, fl. 16 Sept. 1981, *Breteler, Lemmens, Nzabi* 7772 (WAG0449339, WAG0449339, WAG0449340 images!)**; St. Clara**, Tussen ± 50–100 m, Linkerkant, Zij-pod naar St. Clara, sterile, no date, *Breteler* s.n. (WAG044938, image!); **Moyen-Ogooué Province:** “Congo français”. Ogooué (estimated as 0° 41’ 18” S, 10° 13’ 55” E), fl. 1894–95, *Leroy* 23 (PO2093240, PO2093241 images! two sheets).

**Cultivated in Europe exact source unknown**: ex Gabon, probably Sibang, fl. April 2012, leg. *Bogner* 3006 (BR0000019808871, image!).

Those specimens listed above which are sterile, e.g. *Wieringa* 3308 (voucher for DNA studies of the genus, see above), *Wieringa & Haegens* 2710, are only provisionally identified as *P. gabunensis*. It is possible that these specimens might belong to another species of the genus (although unlikely since these specimens were collected at Sibang Arboretum where in recent years only this species of the genus has been collected in flower). Equally they may even represent a species of the genus *Anchomanes*.

**Conservation:** *Pseudohydrosme gabunensis* is possibly extinct at some of its historical locations and is threatened at all of those which remain. At the type location, Sibang, formerly far outside Libreville, at least four gatherings have been made in what is now a small and highly visited forest patch inside Libreville (see notes under *P. buettneri* above). Measured on Google Earth, the forest is approximately a square, c. 470 m N to S, and 420 m W to E, or about 0.25 km^2^ (grid reference: 0° 25’ 56.05”N, 9° 29’ 23.64” E, 49 m alt.). It is now completely surrounded by the dense urban settlement of Libreville which has expanded greatly in the last 60 years. In 1960, at independence, the population of Libreville was 32,000. Since then it has expanded 200-fold to, in 2013, 703,904 (https://en.wikipedia.org/wiki/Libreville, accessed 19 Sept. 2020) and has a vastly greater footprint. Sibang Arboretum, the surviving patch of forest of a once much greater area, is now known as one of the top two tourist destinations in Libreville.

At the Cap Santa Clara location, the Forêt de la Mondah, known since 2012 as the Raponda Walker Arboretum (*Walters et al. 2016*), two collections were made, one in 1981 (see additional collections). Since created as a protected area in 1934, it has been reduced in size, losing 40% of its area in 80 years to habitat clearance and degradation due to its close proximity (c. 15 km) to the metropolis of Libreville which draws upon its trees for timber and firewood (*Walters et al. 2016*). It is not clear whether or not either of the two specimens from St. Clara was from within the current protected area or not.

The species has not been recorded from the Ogouué since it was collected there by Leroy (1894– 1895), despite intensive recent surveys in the lower reaches of the river whence it was probably collected. We have georeferenced the Leroy record from Lambarene since in Leroy’s time this was a trading post n the lower reaches of the river and it is credible that he stopped and collected there, but this is uncertain. Neither has it been recorded in the last century from the Komo at Kango, whence it was collected by Chevalier (26828, P; fl. 2 Oct. 1912). Since this is now on a major transnational route, and on Google Earth shows multiple cleared areas due to development, it is possible that it no longer survives at this location, especially since it has not been recorded here or anywhere near, in such a long time, despite the peak decades of botanical collection in Gabon having been at the end of the 20^th^ century (*Sosef et al., 2005*). *Pseudohydrosme gabunensis* was assessed as Endangered, EN B2ab(ii,iii) by *Lovell & Cheek* (*2020*) since it is or was known from ten specimens at five locations globally, with an area of occupation estimated as 24 km^2^ using the 4 km^2^ cell sizes preferred by IUCN (2012) and the threats detailed above. Threats in the Libreville area have already resulted in the possible global extinction of nine species, including *Pseudohydrosme buettneri* (see under that species, above). The extent of occurrence is calculated as 4,150 km^2^

**Notes**. The location given in the protologue for the type specimen (see above) is similar to that of *Pseudohydrosme buettneri* but more detailed. The Munda is the estuary that forms the eastern edge of the peninsula on which Libreville sits. Tributaries of the Munda drain the Sibang area, one of which may have been known as the Maveli, on the forested banks of which Soyaux recorded collecting the type of *Pseudohydrosme gabunensis*.

The specimens *Leroy* 23 and *Chevalier* 26828 (both P) had been determined as *Amorphophallus* until identified by Bogner (M) as *Pseudohydrosme gabunensis* in Dec. 2012. In contrast, *Wieringa* 4358 (WAG) determined as this species, and cited as such in *Sosef et al*. (*2005*) is in fact an *Amorphophallus*, evident in the larger leaf blade divisions all being acuminate not bifid, and the tuber being described as having the roots from the top (not scattered along the length). Similarly, *Wieringa* 3308 (WAG), correctly cited in *Sosef et al*. (*2005*) as *Pseudohydrosme gabunensis*, was originally collected as an *Anchomanes* until determined by Hetterscheid in April 1996. *Van der Laan* 764 (WAG) had been identified as *Anchomanes nigritianus* Rendle until redetermined by Bogner in Sept. 2012.

*Pseudohydrosme gabunensis* is the most common and widespread member of the genus. However, it is still extremely rare and with a highly restricted range in the wild. It is sought after by private collectors of aroids and live rootstocks and seed attract high prices on the internet. Fortunately, it is found in several large public botanic gardens including in Germany, France, Netherlands, U.K. and U.S.A. We believe that plants are probably not collected from the wild (but this cannot be ruled out), rather they are propagated from those already in cultivation, probably from seed derived from the Netherlands.

The collection reported in *Hetterscheid and Bogner* (*2014*) as from Congo must be treated with great caution. Since it is greatly disjunct (at least c. 300 km) from the known range of this species, it may represent a further new species. If it consists only a leaf and lacks reproductive parts, it may have been confused with an *Anchomanes*. The specimen concerned should be located, studied carefully, and an attempt made to rediscover the source population.

Differences between *Pseudohydrosme gabunensis* and *P. buettneri* are detailed under the last species. There is no doubt that *Pseudohydrosme gabunensis* is much more closely related to *P. ebo* than to *P. buettneri*. However the larger size of the spathes in *P. gabunensis*, their different colour and patterning, the usually bilobed style and bilocular female flowers densely covering the axis, all serve, together with the vegetative characters, to separate it from *P. ebo* (see also table 1 below).

**Table 1.** Characters separating *Pseudohydrosme gabunensis* from *Pseudohydrosme ebo*. Data for *Pseudohydrosme gabunensis* from Bogner (1981:33), Hetterscheid & Bogner (2014:104–113), and live material cultivated at Royal Botanic Gardens, Kew.

|  | *Pseudohydrosme gabunensis* | *Pseudohydrosme ebo* |
| --- | --- | --- |
| Leaf lobe posture and surface | Pendulous, surface bullate (convex between nerves) | Horizontal, surface flat |
| Submarginal nerve in leaf lobes | Absent (veins running to margin) or discontinuous | Continuous |
| Spathe length (cm) | (30–)40–55 | 24–30(–34.5) |
| Spathe tube: colour of outer surface | Uniformly yellow (or white) | Vertical broad pink-brown stripes on white background |
| Spathe blade: colour of inner surface | Yellow or white around margins, central area dark red-purple. Demarcation abrupt | Light reddish brown to pink with pale green veins, gradually slightly darkening in the central area |
| Number of ovary locules & stigma lobes | 2 (–3) | (2–) 3 |
| Female flower density | Covering the rhachis 100% | Covering the rhachis 50–60% |

It is to be hoped that further studies of live plants of *P. ebo* will be possible to determine if, like *P. gabunensis* it also propagates itself apomictically from the root tips.

#### Floral visitors

*Bogner* (*1981*) collected as inferred pollinators two different flies identified as Diptera: Choridae, Sphaeroceridae, and two different beetles identified as Coleoptera: Scaphidiidae, Staphylinidae in association with *Bogner* 664.

#### Reproductive biology

*Hetterscheid & Bogner* (*2014*) working with cultivated plants, report that the female flowering phase is indicated by a faint yet clear lettuce-like scent as the spathe opens, at which time, for 2 days, the receptive stigmas are wet and sticky. After this time, the stigmas turn darker brown, desiccate and are no longer receptive. Individuals are obligate out-crossers. Fruits take up 10 months to mature (*Hetterscheid & Bogner, 2014*).

#### Germination and development

Germination takes 3 weeks to 10 months, producing a single small sagittate, entire leaf from a small rhizome. For several months to two years, new leaves are produced consecutively, usually each larger than its predecessor. From the second leaf onwards, slits may develop in the blade, and within two years the successively produced blades first become divided and finally develop the mature dracontoid pattern (see description). First flowering has occurred in as little as five years from first sowing (*Hetterscheid & Bogner, 2014*). In the wild, time to maturity is likely to take longer due to predation, competition, and likely lower availability of nutrients

**3. Pseudohydrosme ebo** Cheek, *sp. nov*. – Fig. 1–3, 6 & 7.

Differing from *Pseudohydrosme gabunensis* Engl. in the ovaries 3-celled, the stigma conspicuously 3-lobed, very rarely 2-celled/lobed (not: usually 2-celled, 2-lobed, rarely 3-celled/lobed), the female zone of the spadix only c. 50% covered in flowers (not 100% covered), the spathe at anthesis 24– 30(–34.5) cm long, the outer surface dull white with longitudinal brown stripes, inner surface light reddish brown with wide pale green veins (not (30–)40–55 cm long, uniformly white, green or yellow on both surfaces, inner surface bicoloured, the mid-blade area dark purple, separated by a solid line from the marginal white/yellow coloured area). Type: *B.J. Morgan* 25 (holotype K!; isotypes B! MO! YA !), Cameroon, Littoral Region, Yabassi-Yingui, Ebo proposed National Park, fl. Sept. 2010.

Terrestrial herb, to 1.55 m tall. Rhizome cylindric, c. 3 cm diam. obliquely erect to almost parallel to substrate surface, only upper part exposed, surface with transverse ridges (leaf scars) about 2mm deep, 2mm apart. Roots adventitious, thick, fleshy, c. 5 mm diam., scattered along length of rhizome, vegetative apomixis not detected.

Leaf to 1.55 m tall, petiole terete, to 2 cm. diameter at base, green, inconspicuously spotted yellow, mature plants with minute, patent, extremely sparse prickles 0.5 mm long. Blade of youngest seedlings sagittate-elliptic, 5 × 2.5 cm, apex obtuse, basal sinus 1.5 × 1.5 cm, petiole 6–7 cm long. Older seedlings, in successive years with leaves developing first slits and then divisions, becoming triangular in outline with a broad basal sinus. Blade of mature leaves dracontoid, primary division 35–40 × 38–43 cm, pinnatisect, lobes 5–8, dimorphic, larger, mainly distal lobes oblong 12.5–22 × 3.8–6.5 cm, apex acuminate or truncate-bifid (biacuminate), acumen 0.8–1.5 cm long, smaller, mainly proximal lobes ovate c. 8 × 3.5 cm; lateral nerves 6–11 conspicuous on abaxial surface, on each side of the midrib, uniting to form a regular looping submarginal nerve 3–6 mm from the margin, higher order nerves reticulate.

Inflorescence: Cataphylls 4, distichous, light brown, with light green spots, membranous, successively increasing in size from proximal to distal, the outer most triangular-broadly ovate, amplexicaul 3 × 4 cm, the third in succession, long lanceolate-oblong, 12 × 2 cm, the fourth 18–19 × 1.5–4 cm; peduncle 3.5–4.5 × 0.6–0.7 cm, with minute, patent, extremely sparse prickles 0.5 mm long, colour as petiole. Spathe 24–30(–34.5) x 8 cm long basal 1/2–3/4 tubular, funnel-shaped, 1.8–4 cm wide at 2 cm above the peduncle, 6–8 cm wide at 8 cm above the peduncle, and 8–9 cm wide at 15 cm above the peduncle, the distal spathe (blade), half to one third of the total length flaring widely and curving forward, hood-like, shielding the spadix, the apex with a triangular acumen 3–4 × 1 cm. Outer surface of both tube and blade dull white, with pale brown-red ribs running longitudinally along veins from base of tube to mouth of blade. Inner surface of spathe light reddish brown, with wide pale green veins, gradually becoming slightly darker along the midline. Mouth facing horizontally, transversely elliptic, 8–10 cm high, 20–25 cm wide, margin entire. Spadix sessile, cylindrical 50 –85 mm long, 10–18 mm diam. Female zone 24 mm long, 15–18 mm wide, female flowers c. 30 covering about half the surface of the axis, sometimes not contiguous with the male zone the axis then naked for several to 10 mm. Male zone 37–55 mm long, 10–14 mm wide, apex rounded, completely covered in male flowers, sterile appendix absent.

Flowers lacking perigone. Male flowers with c. 5 stamens, stamens free, sessile, prismatic, 5 mm long, in plan-view isodiametric, 5–6 faceted, (1.5–)2 mm diam., apex convex, minutely papillate; anther thecae lateral, four (Fig. 6F) oblong-elliptic, running the length of the stamen, with apical pore (Fig. 6E). Female flowers with ovary globose, 4 mm diam., 3-celled, (Fig. 6I), very rarely 2-celled, style 1–1.5 mm long, 1 mm diam., stigma pale yellow, 0.5 mm thick, 2–2.25 mm wide, strongly 3-lobed (Fig. 7E), lobes with a narrow midline groove, apex rounded. Berry and seed not seen.

**Figure 6.**
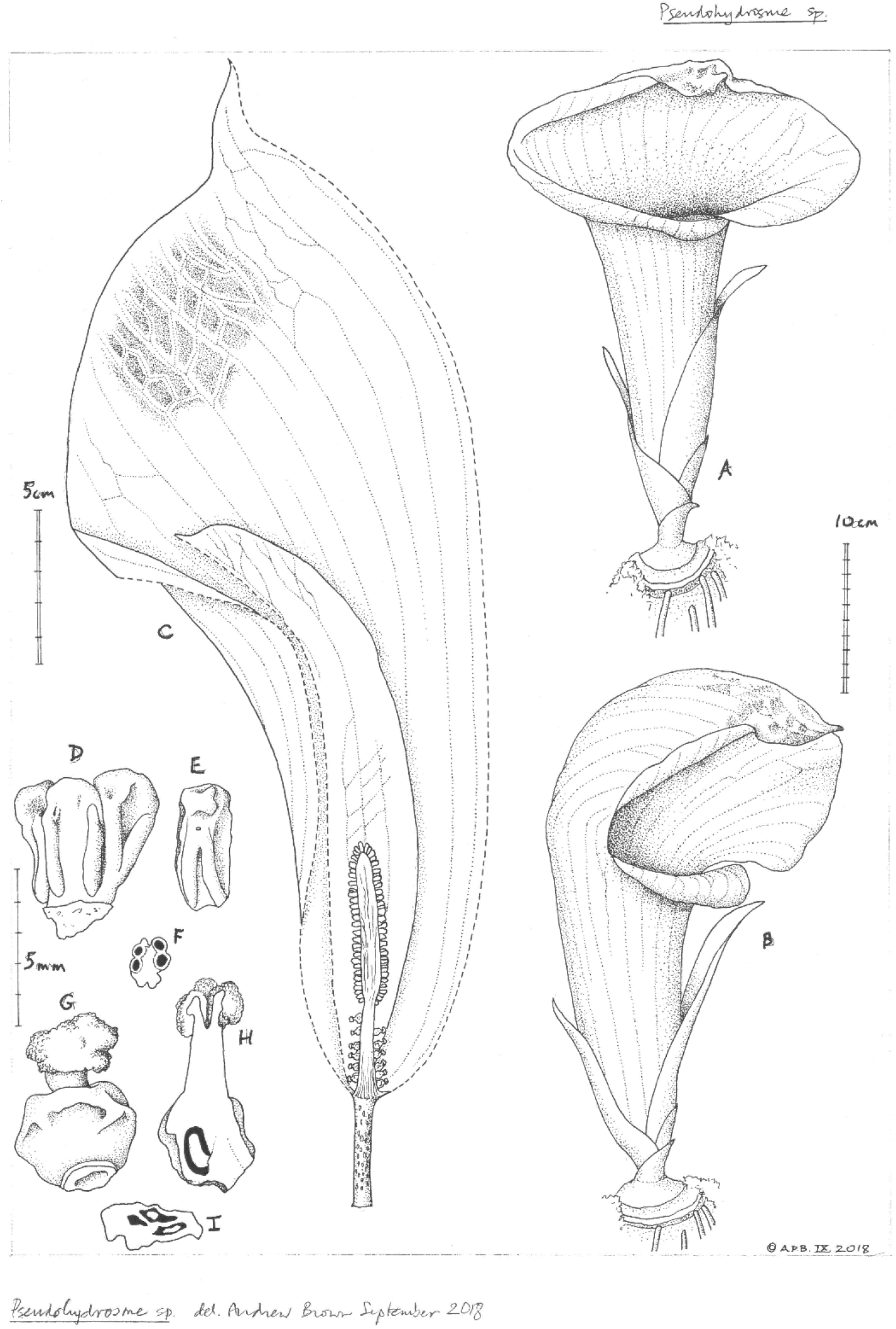
Pseudohydrosme ebo. (*van der Burgt* 1888, K, YA). Drawing of flowering plant. (A & B) habit, rhizome and inflorescences; (C) inflorescence, longitudinal section, showing spadix from base to apex with sparse female flowers, naked axis, male flowers; (D) group of three male flowers, side view; (E) male flower; (F) male flower, transverse section; (G) female flower, entire, side view; (H) female flower, longitudinal section; (I) ovary, transverse section. Drawn by Andrew Brown.

**Figure 7.**
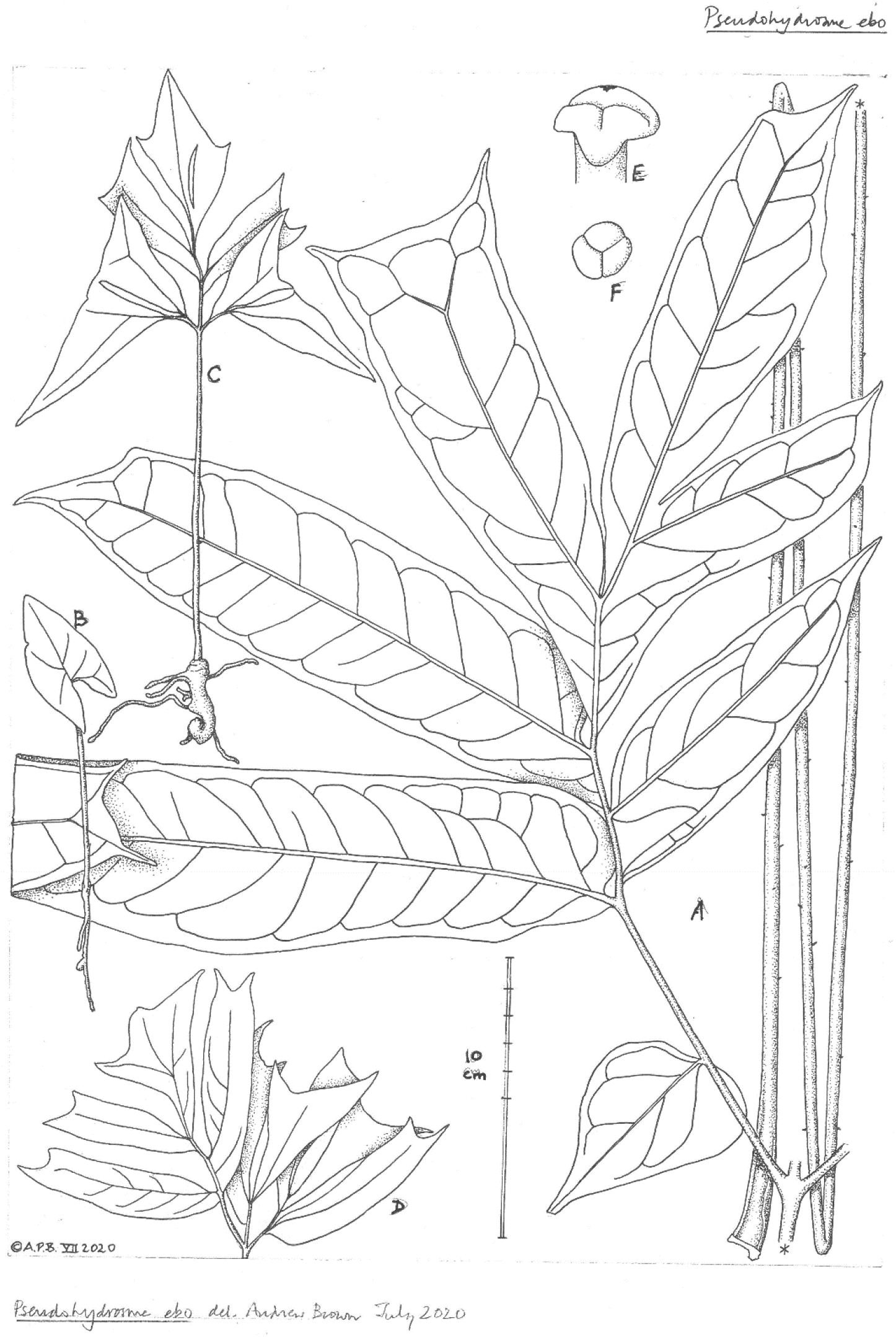
Pseudohydrosme ebo. (A-D from *van der Burgt* 2377, K, YA; E from *Morgan 25*, K,YA). (A) primary division (lateral one of three) of leaf-blade, with petiole folded, behind; (B) seedling (probably in first year); (C) seedling plant (probably in third or later year); (D) seedling leaf-blade (pre-mature); (E & F); trilobed stigma. Drawn by Andrew Brown.

**Distribution & ecology:** Cameroon, Littoral Region, known only from three sites at one location in the Ebo forest near Yabassi-Yingui, in late secondary and intact, undisturbed lowland evergreen forest on ancient basement complex geology, rainfall c. 3 m p.a., drier season October-March; 300– 400 m alt.

**Conservation:** *Pseudohydrosme ebo* is known from only three sites along a section of valley 1.3 km long and only 50–100 mature individuals in total have been seen by the collectors (second and third authors). These sites are along former logging roads which have reverted to forest (third author pers. obs. 2019) as well as intact forest. In the fourteen years since 2006, botanical surveys have been mounted almost annually, at different seasons, over many parts of the formerly proposed National Park of Ebo. About 3000 botanical herbarium specimens have been collected, but despite the species being so spectacular in flower, with individual inflorescences lasting potentially two weeks (if in line with those of *P. gabunensis*), this species has been seen nowhere else in the c. 2000 km^2^ of the Ebo Forest. However, much of this area has not been surveyed during the flowering season of the species, or not surveyed at all for plants. While it is likely that the species will be found at additional sites, there is no doubt that it is genuinely range-restricted. Botanical surveys for conservation management in forest areas neighbouring Ebo resulting in thousands of specimens being collected and identified have failed to find any specimens of *Pseudohydrosme* (*Cheek et al. 1996; Cable & Cheek 1998; Cheek et al. 2000; Harvey et al. 2004; Cheek et al. 2004; Cheek et al. 2010; Harvey et al. 2010*). It is possible that the species is unique to Ebo and truly localised. The area of occupation of *Pseudohydrosme ebo* is estimated as 4 km^2^ using the IUCN preferred cell-size. The extent of occurrence is the same area. In February 2020 it was discovered that moves were in place to convert the forest into two logging concessions (e.g. https://www.globalwildlife.org/blog/ebo-forest-a-stronghold-for-cameroons-wildlife/ and https://blog.resourceshark.com/cameroon-approves-logging-concession-that-will-destroy-ebo-forest-gorilla-habitat/ both accessed 19 Sept. 2020).

This would result in logging tracks that would allow access throughout the forest allowing poachers of rare collectable plants such as *Pseudohydrosme*, and timber extraction would open up the canopy and remove the intact habitat in which *Pseudohydrosme* grows. Additionally, slash and burn agriculture often follows logging trails and would negatively impact the populations of this species. Fortunately the logging concession was suspended due to representations to the President of Cameroon on the global importance of the biodiversity of Ebo (https://www.businesswire.com/news/home/20200817005135/en/Relief-in-the-Forest-Cameroonian-Government-Backtracks-on-the-Ebo-Forest accessed 19 Sept. 2020). However, the forest habitat of this species remains unprotected and threats of logging and conversion of the habitat to plantations remain. *Pseudohydrosme ebo* is therefore here assessed, on the basis of the range size given and threats stated as CR B1+2ab(iii), that is Critically Endangered.

**Additional specimens:** Cameroon, Littoral Region, Ebo proposed National Park, fl. 8 Oct. 2015 *van der Burgt* 1888 (K! YA!); ibid. st. (leaves) 9 Dec. 2019 *van der Burgt* 2377 (K!, MO!, P!, WAG!, YA!).

**Phenology:** flowering in September and early October; leaves early December; fruiting unknown.

**Etymology:** named as a noun in apposition for the forest of Ebo, in Cameroon’s Littoral Region, Yabassi-Yingui Prefecture, to which this spectacular species is globally restricted on current evidence.

**Local names and uses:** none are known.

**Notes:** The discovery of *Pseudohydrosme ebo* is related in the introduction above. Alvarez with van der Burgt, and Ngansop, discovered in Dec. 2019 seedlings of the new species, at three different stages, preserved as *van der Burgt* 2377 sheet ¼ (see Fig 7). Clearly the species at this site is reproducing itself. Associated photographs also show plants of different ages.

*Abwe & Morgan*, (*2008*) and *Cheek et al*. (*2018a*) characterise the Ebo forest, and give overviews of habitats, species, and importance for conservation. Fifty-two globally threatened plant species are currently listed from Ebo on the IUCN Red List website and the number is set to rise rapidly. The discovery of a new species to science at the Ebo forest is not unusual. Since numerous new species have been published from Ebo in recent years. Examples of other species that, like *Pseudohydrosme ebo* appear to be strictly endemic to Ebo on current evidence are: *Ardisia ebo* Cheek (*Cheek & Xanthos, 2012*), *Crateranthus cameroonensis* Cheek & Prance (*Prance & Jongkind, 2015*), *Gilbertiodendron ebo* Burgt & Mackinder (*Burgt et al., 2015*), *Inversodicraea ebo* Cheek (*Cheek et al., 2017*), *Kupeantha ebo* M.Alvarez & Cheek (*Cheek et al., 2018b*), *Palisota ebo* Cheek (*Cheek et al., 2018a*).

Further species described from Ebo have also been found further west, in the Cameroon Highlands, particularly at Mt Kupe and the Bakossi Mts (*Cheek et al., 2004*). Examples are *Myrianthus fosi* Cheek (*Cheek & Osborne, 2010*), *Salacia nigra* Cheek (*Gosline & Cheek, 2014*), *Talbotiella ebo* Mackinder & Wieringa (*Mackinder et al., 2010*)

Additionally, several species formerly thought endemic to Mt Kupe have subsequently been found at Ebo, e.g. *Coffea montekupensis* Stoff. (*Stoffelen et al., 1997*), *Costus kupensis* Maas & H. Maas (*Maas-van der Kamer et al., 2016), Microcos magnifica* Cheek (*Cheek, 2017*), and *Uvariopsis submontana* Kenfack, Gosline & Gereau (*Kenfack et al., 2003*).

Therefore, it is possible that *Pseudohydrosme ebo* might yet also be found in the Cameroon highlands, e.g. at Mt Kupe, further extending westward the known range of the genus. However, this is thought to be only a relatively small possibility given the spectacular nature of this plant, and the high level of survey effort at e.g. Mt Kupe: if it occurred there it is highly likely that it would have been recorded already.

Additional characters separating *Pseudohydrosme ebo* from *P. gabunensis* are show in table 1.

The biogeography of the Cameroonian *Pseudohydrosme ebo* is very different from that of the two Gabonese species of the genus growing c.400 km to the South. The Gabonese species grow on recently deposited, sandy coastal soils. Although the Gabonese species also experience a wet season of about 3 metres of rainfall per annum, it is differently distributed: the dry season in Libreville occurs from June to September inclusive and is colder than the wet season. In contrast at Ebo the geology at the *Pseudohydrosme* location is ancient, highly weathered basement complex, with some ferralitic areas in foothill areas which are inland, c. 100 km from the coast. The wet season (successive months with cumulative rainfall >100 mm) is almost the inverse of at Libreville, falling between March and November and is colder than the dry season (Abwe & Morgan 2008). In addition, the affinities of Ebo as indicated by shared plant species, seems to be with other parts of the Cross-Sanaga biogeographic area, the Cameroon Highlands, rather than with Gabon (see above).

## Discussion

Although indicated as potentially congeneric with *Anchomanes* by *Hetterscheid & Bogner* (*2014*) who cited only the difference in ovary locularity as a basis for maintaining the separation, in fact five other characters support maintaining the separation of these two genera (see table 2 below). Two of these characters were discovered for the first time by those authors. These are 1) the development in the fruit of a pedicel-like stipe and 2) vegetative apomixis from the fleshy roots: producing new plants distant from the parent rhizome. The last character is specifically remarked to be definitively absent from *Anchomanes* species, which have been studied in detail in cultivation (*Hetterscheid, & Bogner 2014*).

**Table 2.** Characters separating *Anchomanes* and *Pseudohydrosme*. Data from *Mayo et al*. (*1997*); *Bogner* (*1981*); *Cusimano et al*. (*2011*); *Hetterscheid & Bogner* (*2014*); this paper.

|  | *Anchomanes* | *Pseudohydrosme* |
| --- | --- | --- |
| Spadix proportions and presentation | Spadix long >1/2–9/10 as long as spathe; conspicuous, projected above the (short) spathe tube | Spadix short, 1/10–1/4 as long as spathe; completely concealed within, and at base of the (long) spathe tube |
| Peduncle | Peduncle exerted far beyond cataphylls; 2–5 x length of spathe | Completely concealed at anthesis by cataphylls; <1/10 length of spathe |
| Ovary locularity and stigma lobe number | 1 | 2 or 3 |
| Berries | Sessile | Stipitate (where known) |
| Laticifers | Present, simple, articulated | Absent |
| Vegetative apomixis via roots | Absent | present |

The description of *Pseudohydrosme ebo* (this paper) necessitates the modification of the circumscription of *Pseudohydrosme* in three respects. Firstly, the leaves have a regular submarginal collective vein, not recorded in *P. gabunensis* and previously a key character used to separate *Amorphophallus* Blume and *Pseudodracontium* N.E.Br. (collective vein present) from *Pseudohydrosme* and *Anchomanes* (continuous submarginal collective vein absent) (Mayo *et al*. 1997:86). Secondly, in *P. ebo* the male and female portions of the inflorescence are not completely contiguous, and the axis is 50% naked in the female portion, while in the other species of *Pseudohydrosme* there is no naked portion and the spadix axis is completely covered in flowers. Thirdly the trilocular ovaries normal in *Pseudohydrosme* ebo are different to those of the other two species which are bilocular, and only very rarely otherwise.

## Conclusions

The discovery of a new species of *Pseudohydrosme* in Cameroon, far from the border with Gabon, is completely unexpected after nearly 130 years in which no additional taxa have been added to the genus. It is also unexpected because one would not predict from the pre-existing data on the genus that such a new species would be so biogeographically and climatically disjunct from its congeners in the Libreville area of Gabon (see under *Pseudohydrosme ebo* above). However examples of even more dramatically unexpected African range extensions have occurred recently such as the westward extension by 2400 km of the genus *Ternstroemia* Mutis ex L.f., of *Talbotiella* Baker by 1400 km, and of the genus *Metarungia* Baden by 1200 km, or in the other direction, eastwards, 1500 km in *Mischogyne* Exell (*Cheek et al. 2019; van der Burgt et al. 2018; Darbyshire et al., 2008*; *Gosline et al*.,*2019* respectively). Such discoveries underline how incomplete our knowledge of the geography of African plant genera remains. Such discoveries also underline the urgency for making such further discoveries while it is still possible since in all but one of the cases given, the range extension resulted from discovery of a new species for science with a narrow geographic range and/or very few individuals, and which face threats to their natural habitat, putting these species at high risk of extinction. About 2000 new species of vascular plant have been discovered each year for the last decade or more. Until species are known to science, they cannot be assessed for their conservation status and the possibility of protecting them is reduced (*Cheek et al., 2020*). Documented extinctions of plant species are increasing, e.g. *Oxygyne triandra* Schltr. of Southwest Region, Cameroon is now known to be globally extinct (*Cheek et al., 2018c*). In some cases species appear to be extinct even before they are known to science, such as *Vepris bali* Cheek, also from the Cross-Sanaga interval in Cameroon (*Cheek et al., 2018d*) and elsewhere, *Nepenthes maximoides* Cheek (*King & Cheek, 2020*). Most of the >800 Cameroonian species in the Red Data Book for the plants of Cameroon are threatened with extinction due to habitat clearance or degradation, especially of forest for small-holder and plantation agriculture following logging (*Onana & Cheek, 2011*). Efforts are now being made to delimit the highest priority areas in Cameroon for plant conservation as Tropical Important Plant Areas (TIPAs) using the revised IPA criteria set out in *Darbyshire et al*. (*2017*). This is intended to help avoid the global extinction of additional endemic species such as *Pseudohydrosme ebo* which will be included in the proposed Ebo Forest IPA.

## Additional Information and Declarations

### Author Contributions

Martin Cheek conceived and designed the experiments, performed the experiments, analyzed the data, wrote the paper, reviewed drafts of the paper. Barthelemy Tchiengue contributed reagents/materials/analysis tools, reviewed drafts of the paper, Xander van der Burgt contributed reagents/materials/analysis tools, reviewed drafts of the paper and produced many of the images.

### Ethics

The following information was supplied relating to ethical approvals (i.e., approving body and any reference numbers):

The fieldwork was approved by the Institutional Review Board of the Royal Botanic Gardens, Kew entitled the Overseas Fieldwork Committee (OFC).

### Field Study Permissions

IRAD-Herbier National du Cameroun sanctioned the field work under a series of Memoranda of Collaboration with the Royal Botanic Gardens, Kew, the most recent signed 4th Sept. 2019, extending until 5^th^ Sept. 2021.

### Data Availability

The following information was supplied regarding data availability:

The specimens on which this manuscript is based are housed in the herbaria for which the standard codes are K and YA. Specimen data and images of type material will be made available on the Kew Herbarium Catalogue at http://apps.kew.org/herbcat/gotoSearchPage.do.

## Acknowledgements

Ekwoge Abwe and Bethan Morgan of San Diego Zoo Global and their team at Ebo Forest Research Project are thanked hugely for expediting our botanical surveys in the Ebo forest of Cameroon over several years. In particular Bethan Morgan is acknowledged for collecting the type specimen of *Pseudohydrosme ebo*.

Janis Shillito is thanked for typing the manuscript. The heads of IRAD (Institute of Research in Agronomic Development)-National Herbarium of Cameroon, Yaounde, successively Jean-Michel Onana, Florence Ngo Ngwe and Eric Nana, are thanked for arranging permits and co-ordinating the co-operation with the Royal Botanic Gardens, Kew. The late Josef Bogner is thanked for conversations on Araceae. Maria Alvarez is thanked for photos of seedlings of *Pseudohydrosme ebo*. Eric Ngansop assisted in the field in Ebo. Marcello Sellaro of the Tropical Nursery, Royal Botanic Gardens, Kew, is thanked for cultivation of and facilitating access to live material of Araceae.

## REFERENCES

Abwe EE, Morgan BJ. 2008. The Ebo Forest: four years of preliminary research and conservation of the Nigeria-Cameroon chimpanzee (Pan troglodytes vellerosus). Pan Africa News 15: 26–29. https://doi.org/10.5134/143494

Amshoff GJH. 1958. Notes on Myrtaceae VII. Myrtaceae of French Equatorial Africa. Acta Botanica Neerlandica 7: 53–58. https://doi:10.1111/j.1438-8677.1958.tb00605.x

Bachman S, Moat J, Hill AW, de la Torre J, Scott B. 2011. Supporting Red List threat assessments with GeoCAT: geospatial conservation assessment tool, in: Smith V, Penev, eds. e-Infrastructures for data publishing in biodiversity science. ZooKeys 150: 117–126. Available from: http://geocat.kew.org/ [accessed 19 July 2020].

Barthlott W, Lauer W, Placke A. 1996. Global distribution of species diversity in vascular plants: towards a world map of phytodiversity. Erkunde 50: 317–328 (with supplement and figure).

Bogner J. 1981. Pseudohydrosme gabunensis Engl. Aroideana 4(1): 31–37.

Bogner J, Petersen G. 2007. The chromosome numbers of the aroid genera. Aroideana 30: 82–90.

Cable S, Cheek M. 1998. The plants of Mt Cameroon, a conservation checklist. Kew: Royal Botanic Gardens.

Cabrera LI, Salazar GA, Chase MW, Mayo SJ, Bogner J, Dávila P. 2008. Phylogenetic relationships of aroids and duckweeds (Araceae) inferred from coding and noncoding plastid DNA. American Journal of Botany 95(9):1153–1165. https://doi:10.3732/ajb.0800073

Cheek M. 1992. Botanical Survey of the Proposed Mabeta-Moliwe Forest Reserve in SW Cameroon: Report on Limbe Gardens Conservation Project. Kew: Royal Botanic Gardens.

Cheek M. 2017. Microcos magnifica (Sparrmanniaceae) a new species of cloudforest tree from Cameroon. PeerJ 5:e4137 https://doi.org/10.7717/peerj.4137

Cheek M, Osborne J. 2010. Myrianthus fosi (Cecropiaceae) a new submontane fruit tree from Cameroon. In: Harvey Y.H., Tchiengue B. & Cheek M. 2010. The plants of the Lebialem Highlands, a conservation checklist: 59–64. Kew: Royal Botanic Gardens.

Cheek M, Xanthos M. 2012. Ardisia ebo sp. nov. (Myrsinaceae) a creeping forest subshrub of Cameroon and Gabon. Kew Bulletin 67: 281–284. https://doi.org/10.1007/s12225-012-9362-8

Cheek M, Alvarez-Agiurre MG, Grall A, Sonké B, Howes M-JR, Larridon L. 2018b. Kupeantha (Coffeeae, Rubiaceae), a new genus from Cameroon and Equatorial Guinea. PLoS ONE 13: 20199324. https://doi.org/10.1371/journal.pone.0199324

Cheek, M, S Cable, FN Hepper, N Ndam & J Watts. 1996. Mapping plant biodiversity on Mt. Cameroon. pp. 110–120 in van der Maesen, van der Burgt & van Medenbach de Rooy (Eds), The Biodiversity of African Plants (Proceedings XIV AETFAT Congress). Kluwer.

Cheek M, Feika A, Lebbie A, Goyder D, Tchiengue B, Sene O, Tchouto P, van der Burgt X. 2017. A synoptic revision of Inversodicraea (Podostemaceae). Blumea 62: 125–156. https://doi.org/10.3767/blumea.2017.62.02.07

Cheek M, Gosline G, Onana J-M. 2018d. Vepris bali (Rutaceae), a new critically endangered (possibly extinct) cloud forest tree species from Bali Ngemba, Cameroon. Willdenowia 48: 285–292. https://doi.org/10.3372/wi.48.48207

Cheek M, Haba PM, Konomou G, van der Burgt, XM. 2019. Ternstroemia guineensis (Ternstroemiaceae), a new endangered, submontane shrub with neotropical affinities, from Kounounkan, Guinea, W. Africa. Willdenowia 49(3): 351–360. https://doi.org/10.3372/wi.49.49306

Cheek M, Harvey Y, Onana JM. 2010. The plants of Dom, Bamenda Highlands, Cameroon, A Conservation Checklist, RBG, Kew, 162pp.

Cheek M, Harvey Y, Onana J-M. 2011. The Plants of Mefou Proposed National Park, Yaoundé, Cameroon, A Conservation Checklist. Kew: Royal Botanic Gardens.

Cheek M, Mackinder B Gosline G, Onana J, Achoundong G. 2001. The phytogeography and flora of western Cameroon and the Cross River-Sanaga River interval. Systematics and Geography of Plants 71: 1097—1100. https://doi.org/10.2307/3668742

Cheek M, Nic Lughadha E, Kirk P, Lindon H, Carretero J, Looney B, Douglas B, Haelewaters D, Gaya E, Llewellyn T, Ainsworth M,Gafforov Y, Hyde K, Crous P, Hughes M, Walker BE, Forzza RC, Wong KM, Niskanen T. 2020. New scientific discoveries: plants and fungi. Plants, People Planet 2:371–388. https://doi.org/10.1002/ppp3.10148

Cheek M, Onana J-M, Pollard BJ. 2000. The Plants of Mount Oku and the Ijim Ridge, Cameroon, a Conservation Checklist. Kew: Royal Botanic Gardens, 220 pp.

Cheek M, Pollard BJ, Darbyshire I, Onana J, Wild C. 2004. The plants of Kupe, Mwanenguba and the Bakossi Mountains, Cameroon. A conservation checklist. Kew: Royal Botanic Gardens.

Cheek M, Prenner G, Tchiengué B, Faden RB. 2018a. Notes on the endemic plant species of the Ebo Forest, Cameroon, and the new, Critically Endangered, Palisota ebo (Commelinaceae). Plant Ecology and Evolution 151(3): 434–441. https://doi.org/10.5091/plecevo.2018.1503

Cheek M, Tsukaya H, Rudall PJ, Suetsugu K. 2018c. Taxonomic monograph of Oxygyne (Thismiaceae), rare achlorophyllous mycoheterotrophs with strongly disjunct distribution. PeerJ 6:e4828 https://doi.org/10.7717/peerj.4828

Cusimano N, Bogner J, Mayo SJ, Boyce PC, Wong M, Hesse M, Hetterscheid WLA, Keating RC, French JC. 2011. Relationships with in the Araceae: comparison of morphological patters with molecular phylogenies. American Journal of Botany 98(4): 1–15. https://doi.org/10.3732/ajb.1000158

Darbyshire I, Anderson S, Asatryan A, et al. 2017. Important Plant Areas: revised selection criteria for a global approach to plant conservation. Biodiversity Conservation 26: 1767–1800. https://doi.org/10.1007/s10531-017-1336-6.

Darbyshire I, Vollesen K, Chapman HM. 2008. A remarkable range disjunction recorded in Metarungia pubinervia (Acanthaceae). Kew Bulletin. 63(4):613–5.

Engler A. 1892. Araceae Africanae I. Botanischer Jahrbucher 15: 447–466.

Engler A. 1911. Pseudohydrosme in: Engler A. Das Pflanzenreich IV 23C(Araceae-Lasioideae): 47–49. Leipzig.

Engler A, Prantl K. 1897. Die natürlichen Pflanzenfamilien nebst ihren Gattungen und wichtigeren Arten: Ergänzungsheft enthaltend die Nachträge zu den Teilen II–IV. W. Engelmann.

Gosline G, Malecot V. 2011. A monograph of Octoknema (Octoknemataceae-Olacaceae s.l.). Kew Bulletin 66(3): 367–404. https://doi.org/10.1007/s12225-011-9293-9

Gosline G, Cheek M, Kami T. 2014. Two new African species of Salacia (Salacioideae,Celastraceae). Blumea 59: 26–32. https://doi.org/10.3767/000651914x682026

Gosline G, Marshall AR & Larridon I. 2019. Revision and new species of the African genus Mischogyne (Annonaceae). Kew Bull 74, 28 (2019). https://doi.org/10.1007/s12225-019-9804-7

Harvey Y, Pollard BJ, Darbyshire I, Onana J, Cheek M. 2004. The plants of Bali Ngemba Forest Reserve, Cameroon. a conservation checklist. Kew: Royal Botanic Gardens.

Harvey YH, Tchiengue B, Cheek M. 2010. The plants of the Lebialem Highlands, a conservation checklist. Kew: Royal Botanic Gardens.

Hay A, Bogner J, Boyce PC. 1994. Nephthytis Schott (Araceae) in Borneo. A new species and a new generic record for Malesia. Novon 4: 65–368. https://doi.org/10.2307/3391445

Hetterscheid W, Bogner J. 2014. Recent observations and cultivation of Pseudohydrosme gabunensis Engl. (Araceae). Aroideana 36: 104–113.

Huynh KL. 1986. Pandanaceae. Flore du Gabon. 28: 3–22.

IPNI. 2020. International Plant Names Index. The Royal Botanic Gardens, Kew, Harvard University Herbaria & Libraries and Australian National Botanic Gardens. Available at http://www.ipni.org (accessed 05 June 2020).

IUCN. 2012. IUCN Red List Categories and Criteria: Version 3.1. Second edition. Gland, Switzerland and Cambridge, UK: IUCN. Available at http://www.iucnredlist.org/ (accessed: 07/2020).

Jacques-Félix H. 1986. Description d’un Tristemma (Melastomataceae) nouveau du Gabon.Bulletin du Muséum national d’Histoire naturelle de Paris, 4e série, section B, Adansonia 8:191–193.

JStor Global Plants. 2020. continuously updated) Available at http://plants.jstor.org/ (accessed 14 June 2020).

Keating RC. 2002. Acoraceae and Araceae. In M. Gregory and D.F. Cutler [eds.], Anatomy of the monocotyledons, vol. 9. Oxford: University Press, Oxford, U.K.

Kenfack D, Gosline G, Gereau, RE, Schatz G. 2003. The genus Uvariopsis in Tropical Africa, with a recombination and one new species from Cameroon. Novon 13: 443–449. https://doi.org/10.2307/3393377

King C, Cheek M. 2020. Nepenthes maximoides (Nepenthaceae) a new, critically endangered (possibly extinct) species in Sect. Alatae from Luzon, Philippines showing striking pitcher convergence with N. maxima (Sect. Regiae) of Indonesia. PeerJ 8:e9899 https://doi.org/10.7717/peerj.9899

Lachenaud O, Stévart T, Ikabanga D, Ngagnia Ndjabouda E, Walters G. 2013. Les forêts littorales de la région de Libreville (Gabon) et leur importance pour la conservation: description d’un nouveau Psychotria (Rubiaceae) endémique. Plant Ecology and Evolution 146(1):68–74. https://doi.org/10.5091/plecevo.2013.744

Lovell R, Cheek M. 2020 (in press). Pseudohydrosme gabunensis. The IUCN Red List of Threatened Species 2020.

Maas-van de Kamer H, Maas PJM, Wieringa JJ, Specht CD. 2016. Monograph of African Costus. Blumea - Biodiversity, Evolution and Biogeography of Plants 61: 280–318. https://doi.org/10.3767/000651916x694445

Mackinder BA, Wieringa JJ, van der Burgt XM. 2010. A revision of the genus Talbotiella Baker f. (Caesalpinioideae: Leguminosae). Kew Bulletin 65: 401–420. https://doi.org/10.1007/s12225-010-9217-0

Mayo SJ, Bogner J, Boyce PC. 1997. The Genera of Araceae. Royal Botanic Gardens, Kew.

Nauheimer LD, Metzler, Renner SS. 2012. Global history of the ancient monocot family Araceae inferred with models accounting for past continental positions and previous ranges based on fossils. New Phytologist.195(4): 938–950.

Omino E. 1996. A contribution to the leaf anatomy and taxonomy of Apocynaceae in Africa: the leaf anatomy of Apocynaceae in East Africa; a monograph of Pleiocarpinae (series of revisions of Apocynaceae 41). Wageningen Agricultural University Papers. 96(1): 1–178.

Onana J, Cheek M. 2011. Red Data Book of the flowering plants of Cameroon, IUCN global assessments. Kew: Royal Botanic Gardens.

Prance GT, Jongkind CCH. 2015. A revision of African Lecythidaceae. Kew Bulletin 70: 6: 13. https://doi.org/10.1007/s12225-014-9547-4

Ruhsam M, Govaerts R, Davis AP. 2008. Nomenclatural changes in preparation for a World Rubiaceae Checklist. Botanical Journal of the Linnean Society. 157(1):115–24.

Stoffelen P, Cheek M, Bridson D, Robbrecht E. 1997. A new species of Coffea (Rubiaceae) and notes on Mt Kupe (Cameroon). Kew Bulletin 52: 989–994. https://doi.org/10.2307/3668527

Szlachetko DL, Olszewski TS. 2001. Orchidacées 3. In: Flore du Cameroun 36. Achoundong G. & Morat P., eds. Paris: Musée National d’Histoire Naturelle; Yaoundé : Herbier National, 666–948.

Sosef MSM, Wieringa JJ, Jongkind CCH, Achoundong G, Azizet Issembé Y, Bedigian D, Van Den Berg RG, Breteler FJ, Cheek M, Degreef J. 2005. Checklist of Gabonese Vascular Plants. Scripta Botanica Belgica 35. Meise: National Botanic Garden of Belgium.

Taton A. 1979. Contribution à l’étude du genre Ardisia Sw. (Myrsinaceae) en Afrique tropicale. Bulletin Du Jardin Botanique National De Belgique / Bulletin Van De National Plantentuin Van België, 49: 81–120. https://doi:10.2307/3667819

Thiers B. 2020. continuously updated. Index Herbariorum: A global directory of public herbaria and associated staff. New York Botanical Garden’s Virtual Herbarium. Available at http://sweetgum.nybg.org/ih/ (accessed June 2020).

Thistleton-Dyer. 1901. Flora of Tropical Africa 8. London: L. Reeve & Co.

van der Burgt XM, Mackinder BA, Wieringa JJ et al. 2015. The Gilbertiodendron ogoouense species complex (Leguminosae: Caesalpinioideae), Central Africa. Kew Bulletin 70: 29. https://doi.org/10.1007/s12225-015-9579-4

van der Burgt XM, Molmou D, Diallo A. et al. 2018. Talbotiella cheekii (Leguminosae: Detarioideae), a new tree species from Guinea. Kew Bulletin 73, 26 https://doi.org/10.1007/s12225-018-9755-4

Walters G, Ndjabounda EN, Ikabanga D, Biteau JP, Hymas O, White LJ, Obiang AM, Ondo PN, Jeffery KJ, Lachenaud O, Stevart T. 2016. Peri-urban conservation in the Mondah forest of Libreville, Gabon: Red List assessments of endemic plant species, and avoiding protected area downsizing. Oryx 50(3):419–30. https://doi.org/10.1017/s0030605315000204

